# Predictive value of preclinical models for CAR-T cell therapy clinical trials: a systematic review and meta-analysis

**DOI:** 10.1101/2024.12.15.628103

**Authors:** David Andreu-Sanz, Lisa Gregor, Emanuele Carlini, Daniele Scarcella, Carsten Marr, Sebastian Kobold

## Abstract

Experimental mouse models are indispensable for the preclinical development of cancer immunotherapies, whereby complex interactions in the tumor microenvironment (TME) can be somewhat replicated. Despite the availability of diverse models, their predictive capacity for clinical outcomes remains largely unknown, posing a hurdle in the translation from preclinical to clinical success. This study systematically reviews and meta-analyzes clinical trials of chimeric antigen receptor (CAR-) T cell monotherapies with their corresponding preclinical studies. Adhering to PRISMA guidelines, a comprehensive search of PubMed and ClinicalTrials.gov was conducted, identifying 422 clinical trials and 3157 preclinical studies. From these, 105 clinical trials and 180 preclinical studies, accounting for 44 and 131 distinct CAR constructs, respectively, were included.

Patientś responses varied based on the target antigen, expectedly with higher efficacy and toxicity rates in hematological cancers. Preclinical data analysis revealed homogenous and antigen-independent efficacy rates. Our analysis revealed that only 4 % (n = 12) of mouse studies used syngeneic models, highlighting their scarcity in research. Three logistic regression models were trained on CAR structures, tumor entities, and experimental settings to predict treatment outcomes. While the logistic regression model accurately predicted clinical outcomes based on clinical or preclinical features (Macro F1 and AUC > 0.8), it failed in predicting preclinical outcomes from preclinical features (Macro F1 < 0.5, AUC < 0.6), indicating that preclinical studies may be influenced by experimental factors not accounted for in the model.

These findings underscore the need for better understanding the experimental factors enhancing the predictive accuracy of mouse models in preclinical settings.

## Introduction

In cancer immunotherapy, CAR-T cells are engineered with a synthetic receptor targeting tumor antigens, leading to T cell activation and subsequent cancer cell killing^1,2^. Approved therapies including Kymriah®, Yescarta® and Tecartus® have shown short and long-term success in hematological cancers^3^, where tumor cells are readily available in the bloodstream or bone marrow. Nevertheless, a substantial proportion of patients will still not benefit or only transiently respond^4-6^, while experiencing severe adverse events (AE)^7-9^. Conversely, solid tumors possess physical barriers compromising accessibility of therapeutic cells, resulting in lower response rates^10-11^. Despite promising results for targets like human epithelial growth factor 2 (HER2)^12-15^, mesothelin^16-18^, and disialoganglioside (GD2)^19,20^, clinical efficacy in solid tumors remains limited. We and others have identified access to cancer tissue, limited tumor cell recognition and immune suppression as key resistance mechanisms contributing to clinical inefficacy^4,21,22^.

Overall, translation of preclinical drugs to market authorization stands at a dramatic 1 in 10.000, with a phase I clinical trialś success rate below 5 %^23^. Enhancing the predictive capacity of preclinical studies regarding their therapeutic window and future clinical efficacy is essential for increasing productivity of drug development processes. For CAR-T cells, regulatory agencies mandate preclinical animal testing prior to clinical development^23^ and beside canine or non-human primate models, the majority of preclinical CAR-T cell development relies on mouse models^24,25^, making these a key stepping stone for decision-making during drug development (Table 1).

**Table 1:** Overview of different mouse models used in preclinical cancer immunology research including their advantages and disadvantages.

|  | Advantages | Disadvantages |
| --- | --- | --- |
| Syngeneic<br>(transplanted<br>tumor cell lines) | • Cost efficiency and time saving model<br>• Possible to study on-target off-tumor effects<br>• Efficacy study concomitantly with native immune system<br>• Possibility of orthotopic implantation | • Very limited tumor heterogeneity<br>• Limited modeling of TME<br>• Limited comparison to clinical (human) setting<br>• Lack of human binder-mouse antigen cross-reactivity<br>• Artificial tumor location if implemented subcutaneously |
| Syngeneic<br>(spontaneous<br>induced cancer<br>development) | • Cost efficient model<br>• More realistic model of tumor development and progression<br>• Possible to study on-target off-tumor effects<br>• Efficacy study concomitantly with native immune system<br>• Potential orthotopic study of tumor model | • Variation in spontaneous mutation rates thus limited comparability<br>• Variation in onset and location of tumor development<br>• Time-consuming model due to tumor development<br>• Limited comparison to clinical (human) setting |
| Transgenic<br>models | • Controlled genetic manipulation of the tumor<br>• Close modeling of human tumors<br>• Targeted insights regarding specific genes possible | • High maintenance due to advanced gene editing techniques<br>• Time-consuming model due to tumor development<br>• Possibility of unintended phenotypes to occur<br>• |
| Xenograft models | • Easy implementation and time saving model<br>• Allows experimentation with human cancer cell lines and human T cells<br>• Suitable to study human tumor biology | • Use of human cell lines limits the tumor heterogeneity<br>• Graft vs. Host disease<br>• High costs of immunodeficient mice<br>• Immunocompromised host limits immune response studies |
| Patient-derived xenograft (PDX) | • Use of patient specific tumor cells increases clinical relevance<br>• Maintains tumor heterogeneity<br>• Can reflect patient-specific treatment responses (at least with regards to tumor cell directed effects) | • GvHD<br>• Limited access to suitable patient samples<br>• High costs and time-consuming model |

Syngeneic models using spontaneous, induced, or transplantable tumors in mice with C57BL/6 or BALB/c backgrounds allow the study of adoptive transfer alongside a fully-competent host immune system^26-28^. Despite their genomic homogeneity, these models face clinical comparability issues due to their artificial TME^26,27,29,30^. Subcutaneous implantation simplifies tumor measurements, but lacks the complexity of natural tumor growth^31^. Orthotopic transplantation on the other hand may offer a more accurate tumor reflection, but requires training and complex surgical procedures^32-34^.

Human xenograft models consist of transplanting human tumor and immune cells into immunocompromised mice such as NOD/SCID/IL2Rγc-KO (NSG) and enable the study of human tumor-immune cell interactions^35,36^. Immunodeficient mice pose challenges including graft-versus-host disease (GvHD), incomplete human immunity^27,37-39^, and the absence of human stromal cells, which impairs CAR-T cell preservation and limits the study of tumor-supporting stroma in solid tumor therapies^27,37,38^. Patient-derived xenografts (PDX) replicate tumor cell-intrinsic features into immunodeficient mice, but typically have slow growing tumors and lack human hematopoiesis, complicating immunotherapy evaluation^40-42^.

Despite their disadvantages, preclinical mouse models remain crucial for advancing therapies up to clinical testing. Proactive and comprehensive assessment of potential side effects and toxicities to reliably predict clinical efficacy and safety is imperative^8,43,44^.

Promising preclinical results, especially in solid tumors, often fail to translate into strong clinical responses. Limited understanding of the accuracy of these results in forecasting clinical responses in patients, coupled with a restricted therapeutic evaluation in a single model, could explain this gap. To gain a better understanding of the performance of preclinical models in CAR-T cell therapy development, we set out to perform a meta-analysis of all available clinical trials investigating CAR-T cell monotherapies together with their preclinical workup in mouse models. Using machine learning models, we sought to predict clinical trial outcomes based on preclinical data and identify potential factors that allow such forecasts.

## Materials and Methods

### Information sources, search strategy, and data collection process

The clinical trial and preclinical records were sourced from PubMed and ClinicalTrials.gov until December 1^st^, 2023, employing specific search criteria (Supplementary methods). These included all publications in English, excluding reviews, systematic reviews, meta-analyses, and retrospective studies. To prevent biases in the assessment, each entry was evaluated in all its aspects, including its fit with regard to inclusion/exclusion criteria and data extraction, by at least two reviewers (D.A.S., L.G., E.C.) independently.

### Eligibility criteria and selection process

Clinical trial entries were excluded if they fulfilled at least one of the following criteria: Non-cancer related, Not CAR-T cell therapy, Follow-up study, Retrospective study, Results not available, Terminated for non-scientific reasons, Not evaluating efficacy, Combination therapy, Case-report, Reported somewhere else, Incomplete data. Conversely, preclinical entries were excluded if fulfilling any of the following criteria: Not original research article, Non-cancer related, Clinical study, Not CAR-T cell therapy, Non-classical CAR setting, Target not in clinical setting, Not about CAR efficacy, No animal models, Follow-up studies, Retrospective study, Combination therapy, Incomplete data, Studying irrelevant variables in the experimental set-up. An additional clarification of the exclusion criteria considered by the investigators as the most interpretative is provided in the supplementary methods.

### Analysis of clinical trials

For included clinical studies, pre-established variables were collected (Table 2): the nature of the tumor entity and target antigen; the full structure of the CAR: single-chain variable fragment (scFv), transmembrane, intracellular domains; the administered chemotherapeutic lymphodepleting regimen; the use of hematopoietic stem cell transplantation before or after CAR-T cell therapy for all patients; and the number of participants belonging to the efficacy, and the safety assessments of the CAR-T cell therapy of interest. Evaluating the efficacy consisted in dividing the patient population into four categories of overall response, namely (i) progressive disease (PD), (ii) stable disease (SD), (iii) partial response (PR), and (iv) complete response (CR). If not clearly stated, but discernible by the published data, the latest time point of disease assessment was considered. For the safety assessment of each CAR-T cell therapy, all serious and less severe AE were reported. When side effects were evaluated at multiple time points, the earliest time point after infusion was considered. The severity of all AE was classified into mild (Grade 1-2) or severe (Grade 3-4), as stated by the investigators. The list of AE included (i) cytokine release syndrome (CRS), (ii) immune effector cell-associated neurotoxicity syndrome (ICANS), (iii) graft versus host disease (GvHD), (iv) hematotoxicity (distinguished between anemia, leukopenia, lymphocytopenia, thrombocytopenia and neutropenia), and (v) death.

**Table 2:** Summary of all variables collected for clinical trials analyzed.

|  |  |
| --- | --- |
| CAR Structure | CAR generation |
|  | scFv source |
|  | Transmembrane Domain |
|  | Costimulatory domain |
|  | Signaling domain |
| Additional Therapy | Preconditioning therapy |
|  | Stem cell transplantation |
| Response | Progressive Disease |
|  | Stable Disease |
|  | Partial Response |
|  | Complete Response |
| Adverse Events (AE) | GvHD (Grade 1-5) |
|  | CRS (Grade 1-5) |
|  | ICANS (Grade 1-5) |
|  | Treatment-related deaths |
|  | Anemia (Grade 1-5) |
|  | Thrombocytopenia (Grade 1-5) |
|  | Lymphopenia (Grade 1-5) |
|  | Leukopenia (Grade 1-5) |
|  | Neutropenia (Grade 1-5) |

### Analysis of preclinical publications

All approved preclinical studies underwent thorough review and data extraction, with pre-established variables collected. Multiple entries were created for different mouse models or CAR therapies within a single publication (Table 3). Variables studied included: the nature of the tumor entity, the origin and expression density of the target antigen; the mouse strain; the route of tumor cell inoculation; the full structure of the CAR; the use of a preconditioning regimen; and the number of all mice in the study treated with the CAR-T cell therapy of interest. The overall response of each treatment group was determined by the average of all reported responses within that group. The safety assessment encompassed all reported cases of AE such as (i) relapse, (ii) weight loss, (iii) CRS, (iv) ICANS, (v) GvHD, and (vi) death of experimental animals due to therapy side effects.

**Table 3:** Overview of all variables collected for all preclinical publications analyzed.

|  |  |
| --- | --- |
| Mouse Model | Disease entity category |
|  | Source of the cancer cell line |
|  | Type of antigen |
|  | Antigen overexpression |
|  | Any further genetic modification |
|  | Injection site of tumor cell line |
|  | Mouse strain |
|  | Mouse type |
|  | Preconditioning therapy |
|  | Rechallenge |
| CAR Structure | CAR generation |
|  | scFv source |
|  | Transmembrane domain |
|  | Costimulatory domain |
| Response | Tumor decrease |
|  | Tumor stable |
|  | Tumor clearance |
|  | Relapse |
| Adverse Events (AE) | Weight decrease |
|  | CRS |
|  | GvHD |
|  | Treatment-related deaths |

### Machine learning-guided analysis

Our analysis used both the preclinical (n = 303 experimental entries) and clinical (n = 105 clinical trials) CAR-T cell datasets. The response variables were binarized for logistic regression (Figure 4): In preclinical data individual mice were categorized into ‘No response’ vs. ‘Response’, while in clinical data, an overall response rate (ORR) cutoff of 0.25 was used (Individuals with an ORR < 0.25: ‘No response’; those with ORR >= 0.25: ‘Response’). Three logistic regression models were developed: Model A was trained and validated on preclinical data with 14 features; hyperparameters were optimized using grid search with 5-fold cross-validation (Supplementary methods). Model B was trained on preclinical data and validated on clinical data, utilizing shared 7 features (’Solid or Hematologic tumors’, ‘scFv source’, ‘Target’, ‘CAR generation’, ‘TM domain’, ‘Preconditioning’, ‘Costimulatory domain’). Model C was trained and validated on clinical data with 7 features. Performance metrics included AUC and macro F1 score. Feature importance was assessed through logistic regression coefficients. Datasets were further divided into solid and hematological tumors, with separate models trained and evaluated for each subset. A Lasso linear regression model D was applied to predict the continuous ORR in clinical data, with feature importance determined by non-zero coefficients (Supplementary methods). Analysis was performed in a conda environment using Python 3.10.13, scikit-learn 1.3.2, and seaborn 0.13.0.

## Results

### Literature review and data collection

This study sought to bridge the gap between preclinical and clinical data in CAR-T cell therapy, not only by evaluating the clinical trial landscape and its respective preclinical publications, but also by using a logistic regression model to test whether preclinical studies can predict clinical treatment outcomes. A systematic review of clinical and preclinical CAR-T cell studies was conducted following PRISMA guidelines^45^ (Figure 1). At all times, two researchers independently screened the results, identifying 422 clinical and 3157 preclinical articles, and excluded articles due to combinatorial therapies, mislabeling or lacking data. 105 clinical and 180 preclinical studies met inclusion criteria, with no overlap of patients or animals occurring between studies.

**Figure 1:**
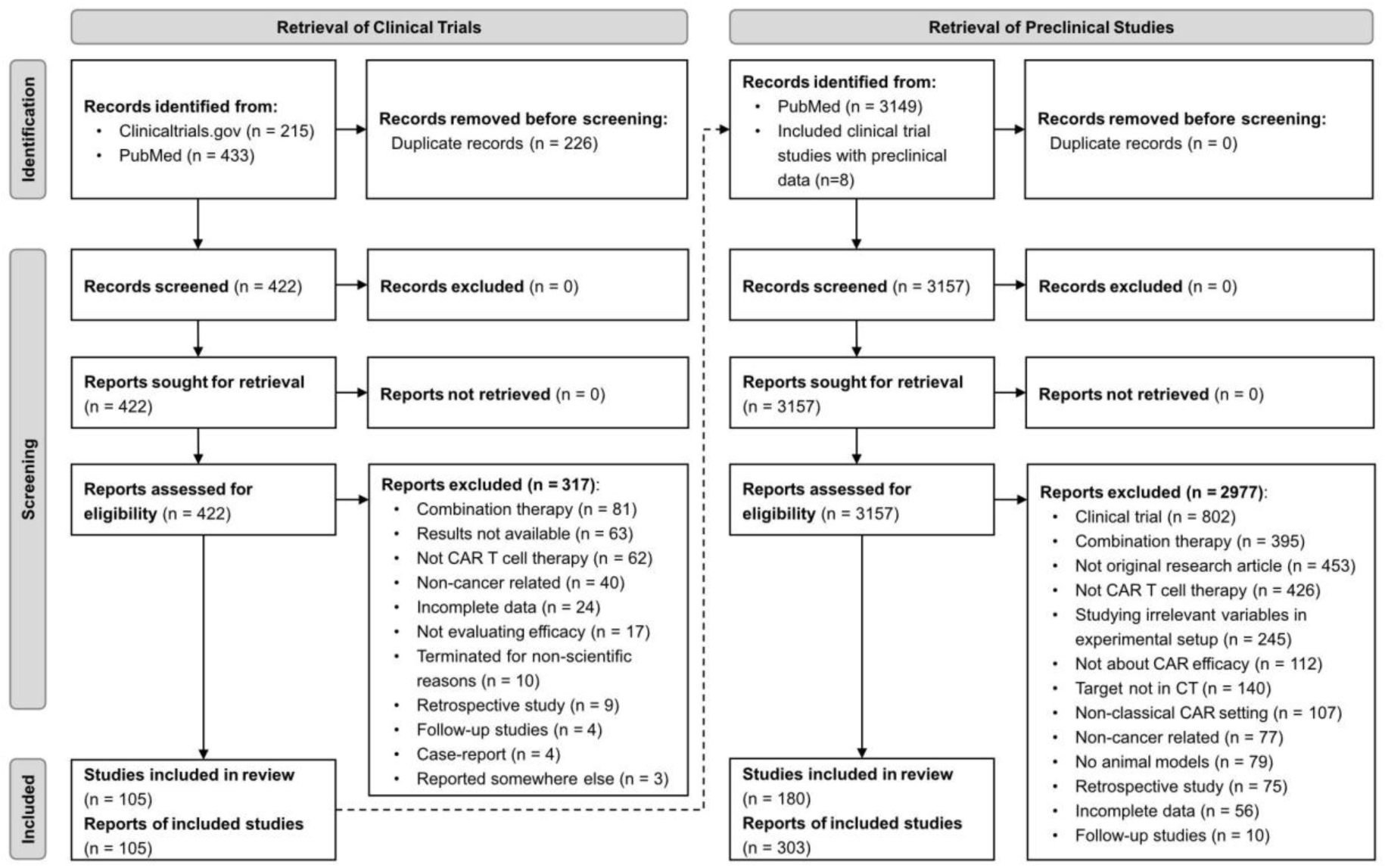
Overview of literature review and data collection process. Literature search performed until December 1^st^ 2023 on PRISMA 2020 led to 105 clinical trials included in our study. Using the target antigens from the included clinical trials (dotted arrow), 303 relevant preclinical studies employing the same target antigen were identified and included in our analysis.

### Increased therapeutic efficacy and higher rates of side effects in clinical trials of hematological tumors

Searching PubMed, all clinical trials involving CAR-T cell therapy were retrieved until December 1st 2023, of which 105 clinical trials met inclusion criteria. 86 trials focused on hematological cancers, primarily employing anti-CD19 (n = 53) and anti-B cell maturation antigen (BCMA) (n = 16) CAR-T cells (Figure 2A). The meta-analysis included 3312 patients with hematological cancers and 184 with solid cancers (Figure 2B, Figure S1B), with most T cell products using murine-derived scFvs, second-generation CAR structures with 4-1BB or CD28 co-stimulatory domains (91 %) (Figure 2C). As expected, hematological cancer trials showed higher ORR than those for solid tumors, with only one trial involving anti-GD2 or anti-mesothelin CAR-T cells showing notable efficacy, respectively (Figure 2D, E). Interestingly, preconditioning therapy, such as chemotherapy or irradiation, was used in 90 % of hematological cancer trials and in 37 % of all solid cancer trials (Figure S1A). Expectedly, higher clinical responses in hematological cancers were strongly correlated with an increased rate of side effects, including ICANS and CRS (Figure 2F-H). GvHD was rare, mainly occurring in patients treated with murine scFv CARs (Figure 2I). Hematotoxicity symptoms were less common in solid cancer patients (Figure 2J-N), partly explainable by the limited use of preconditioning therapy.

**Figure 2:**
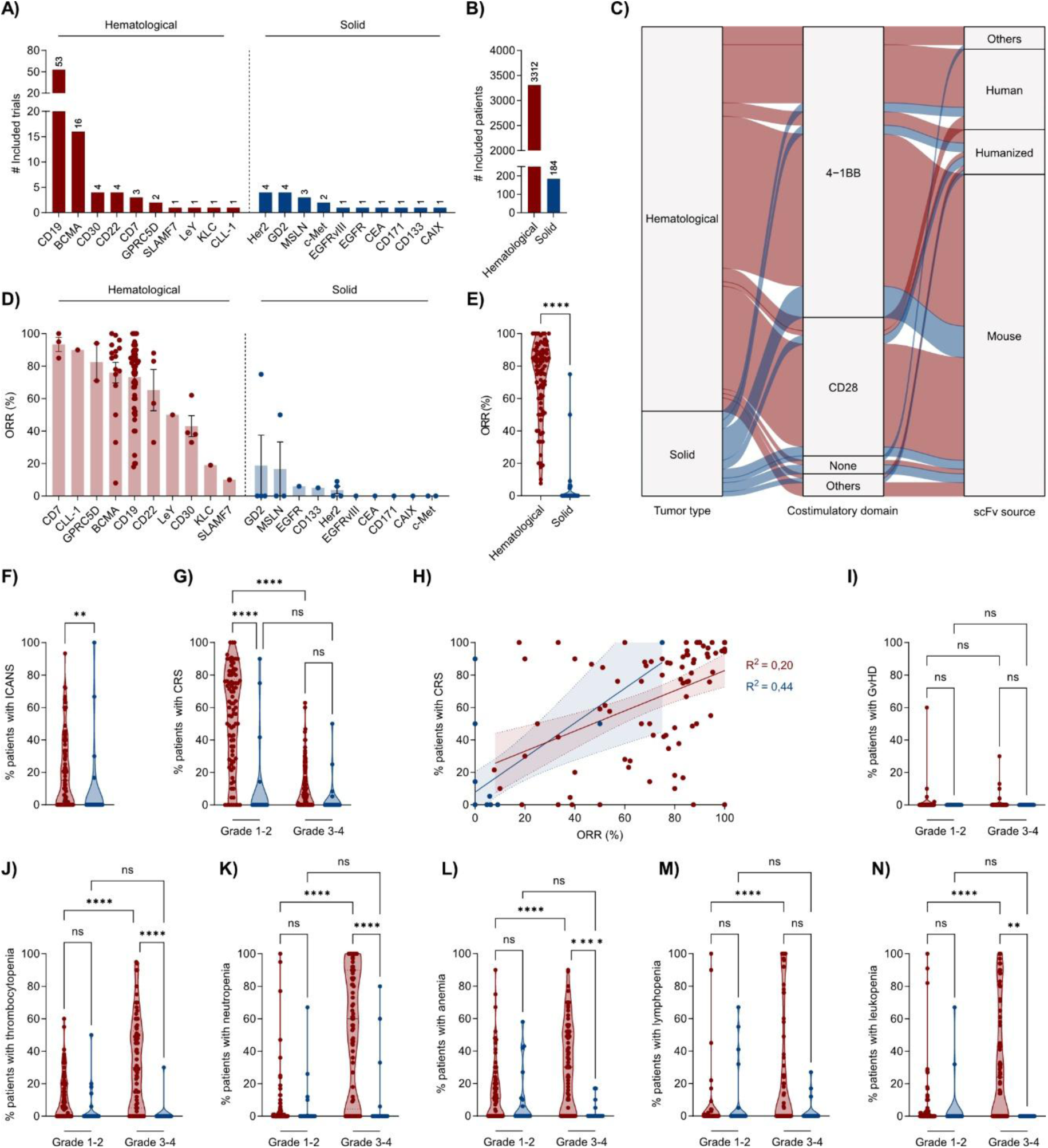
Hematological tumors are associated with higher response rate and toxicity than solid tumors in clinical CAR-T cell trials. (A) Number of clinical trials analyzed for each included target antigen. (B) Total number of included patients for hematological and solid cancer clinical trials. (C) Detailed information regarding costimulatory domain of CAR construct, as well as source of scFv. (D) ORR separated by target antigen. (E) ORR of all clinical trials for hematological and solid tumor entities. (F) Percentage of patients experiencing ICANS in hematological and solid tumor entities. (G) Percentage of patients experiencing CRS in hematological and solid tumor entities. (H) Correlation between ORR and occurrence of side effects in terms of CRS in patients. (I) Fraction of patients experiencing GvHD. Distribution of patients experiencing (J) thrombocytopenia, (K) neutropenia, (L) anemia, (M) lymphopenia or (N) leukopenia.

### Preclinical mouse studies of CAR-T cell therapy in hematological and solid cancers

Preclinical publications on CAR-T cell therapy targeting antigens from prior clinical trials were retrieved from PubMed by December 1st, 2023. Most studies focused on anti-CD19 (n = 77), anti-mesothelin (n = 53), or anti-HER2 CAR-T cells (n = 37) (Figure 3A). Most B cell acute lymphoblastic leukemia models (85 %) were utilizing anti-CD19 CAR-T cells, while CAR targets for solid tumors covered a comparably wider range of malignancies (Figure 3B). The number of mice (hematological n = 1121, solid n = 1311) used for *in vivo* validation was similar for both tumor types (Figure 3C). Partial responses, including tumor reduction and slower growth, were most commonly reported in immunodeficient mouse models, namely in 65,3 % of solid tumors and 72,7 % of hematological tumor entities (Figure 3D-E). Complete responses were more frequent in the small number of reported immunocompetent models, namely 47,6 % for solid tumors and 45,5 % for hematological tumors (Figure 3F). Expectedly, most CAR-T cells tested (69 %), used murine scFvs with 4-1BB or CD28 as co-stimulatory domains. CAR-T cells targeting hematological tumors favored 4-1BB co-stimulation, while solid tumor CARs leaned toward CD28 (Figure 3G). In general, preclinical toxicity was reported in only 4 % of the publications studied, with few studies noting issues like weight loss, CRS, ICANS, GvHD, or lethal AE (Figure S2A-F). Only a small fraction (n = 8) were fully syngeneic, using murine antigens and tumor cells in immunocompetent mice (Figure 3H, Figure S2G-H). Only 3 % of all reported solid tumor experiments and 6,6 % of all hematological tumors used PDX models, which predominantly demonstrated partial responses (75 % for solid tumors and 77 % for hematological tumors) (Figure S2I-J). This highlights a bias towards an increased use of immunodeficient models in preclinical CAR-T cell testing, with easier to achieve responses and minimal safety evaluation.

**Figure 3:**
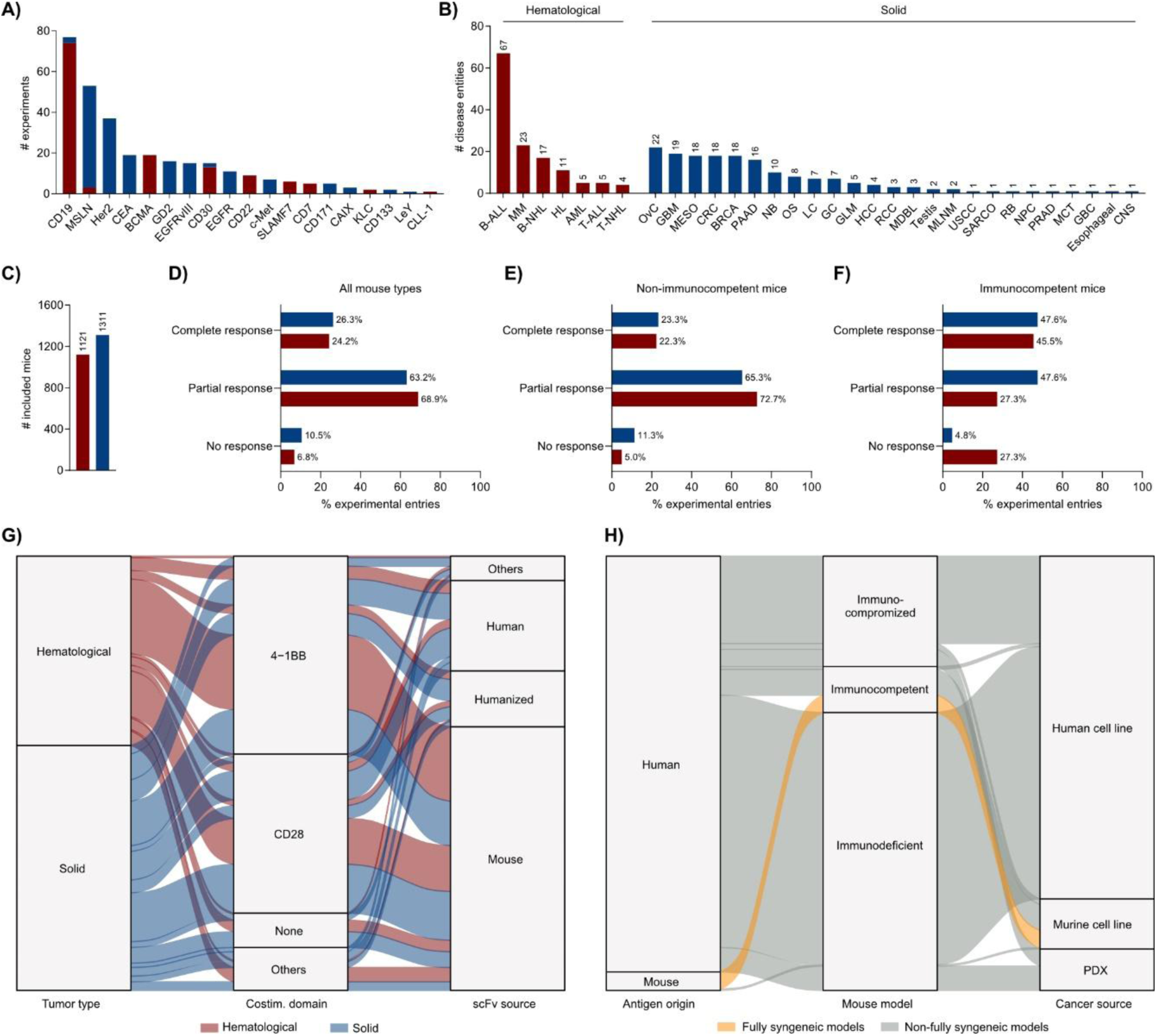
Preclinical mouse studies of CAR-T cell therapy in hematological and solid cancers are primarily of immunodeficient nature. (A) Overall number of preclinical entries per CAR target for either hematological or solid tumors. (B) Overall number of preclinical entries per disease entity for either hematological or solid tumors. (C) Sum of all mice belonging to CAR treatment groups for either category of tumors. (D) Overall responses as for tumor clearance, decrease, slower growth or progression for all mice, non-immunocompetent mice (E) and immunocompetent mice (F) for either type of tumor. (G) Alluvial plot displaying the different proportions of preclinical studies according to the scFv and co-stimulatory domain of their CAR molecule for either type of tumor. (F) Alluvial plot highlighting, in yellow, the proportion of preclinical studies employing fully syngeneic mouse models for in vivo CAR-T cell investigation. Abbreviations: B-ALL, B-cell acute lymphoblastic leukemia; MM, multiple myeloma; B-NHL, B-cell non-Hodgkin lymphoma; HL, Hodgkin lymphoma; T-ALL, T-cell acute lymphoblastic leukemia; AML, acute myeloid leukemia; T-NHL, T-cell non-Hodgkin lymphoma; OVC, ovarian cancer; GBM, glioblastoma; BRCA, breast adenocarcinoma; CRC, colorectal carcinoma; MESO, mesothelioma; PAAD, pancreatic adenocarcinoma; NB, neuroblastoma; OS, osteosarcoma; GC, gastric cancer; LC, lung cancer; GLM, glioma; HCC, hepatocellular carcinoma; MDBL, medulloblastoma; RCC, renal cell carcinoma; MLNM, melanoma; TC, testicular cancer; CNS, central nervous system tumors; ESO, oesophageal cancer; GBC, gallbladder cancer; NPC, nasopharyngeal cancer; MCT, mast cell tumor; PRAD, prostate adenocarcinoma; RB, retinoblastoma; SARCO, sarcoma; USCC, serous carcinoma of the uterine cervix.

### Preclinical and clinical data can predict clinical treatment outcome using a logistic regression model

To evaluate the predictive value of preclinical models in CAR-T cell trials, a comparative machine learning analysis was conducted. Three logistic regression models (A, B, and C) and a linear regression model (Model D) were deployed using different sets of training and testing data (Figure 4A): Models A to C were trained on subsets, i.e. (i) ‘all tumors’, (ii) ‘hematologic tumors’ only, and (iii) ‘solid tumors’ only. Due to the small sample size and label imbalance in the clinical solid tumor subset (17 non-responders vs. 2 responders), only Model A was trained and tested on the preclinical solid tumor data. When both hematological and solid tumor types (‘all tumors’) were included, Models B and C achieved higher predictive power than Model A (Figure 4B-C), which showed no performance beyond random guessing (Macro F1 ∼ 0.4 and AUC ∼ 0.5, respectively). To exploit the continuous nature of the ORR in clinical studies, we also trained a regularized linear regression model D to assess the predictive value of clinical features. Although this model demonstrated poor overall performance (mean R²: 0.51), it correctly predicted a lower overall response rate (ORR) for solid tumors (Figure 4E), with the largest negative weight assigned to the ‘Solid’ tumor type (Figure 4F). ‘Tumor type’ (‘Solid’ or ‘Hematological’) emerged as the most relevant discriminator in both classification models B and C (Figure 4D) and the regression Model D (Figure 4F), indicating that preclinical and clinical data can predict clinical outcome when tumor type information is included. When Model C is trained and tested on the ‘hematologic tumor’ subset, its performance is still beyond random guessing (mean F1 = 0.55 ± 0.14, Figure 4B). ‘TM Domain’ is identified as the most predictive feature, followed by ‘Preconditioning’ and the ‘Costimulatory domain’ (Figure 4D).

**Figure 4:**
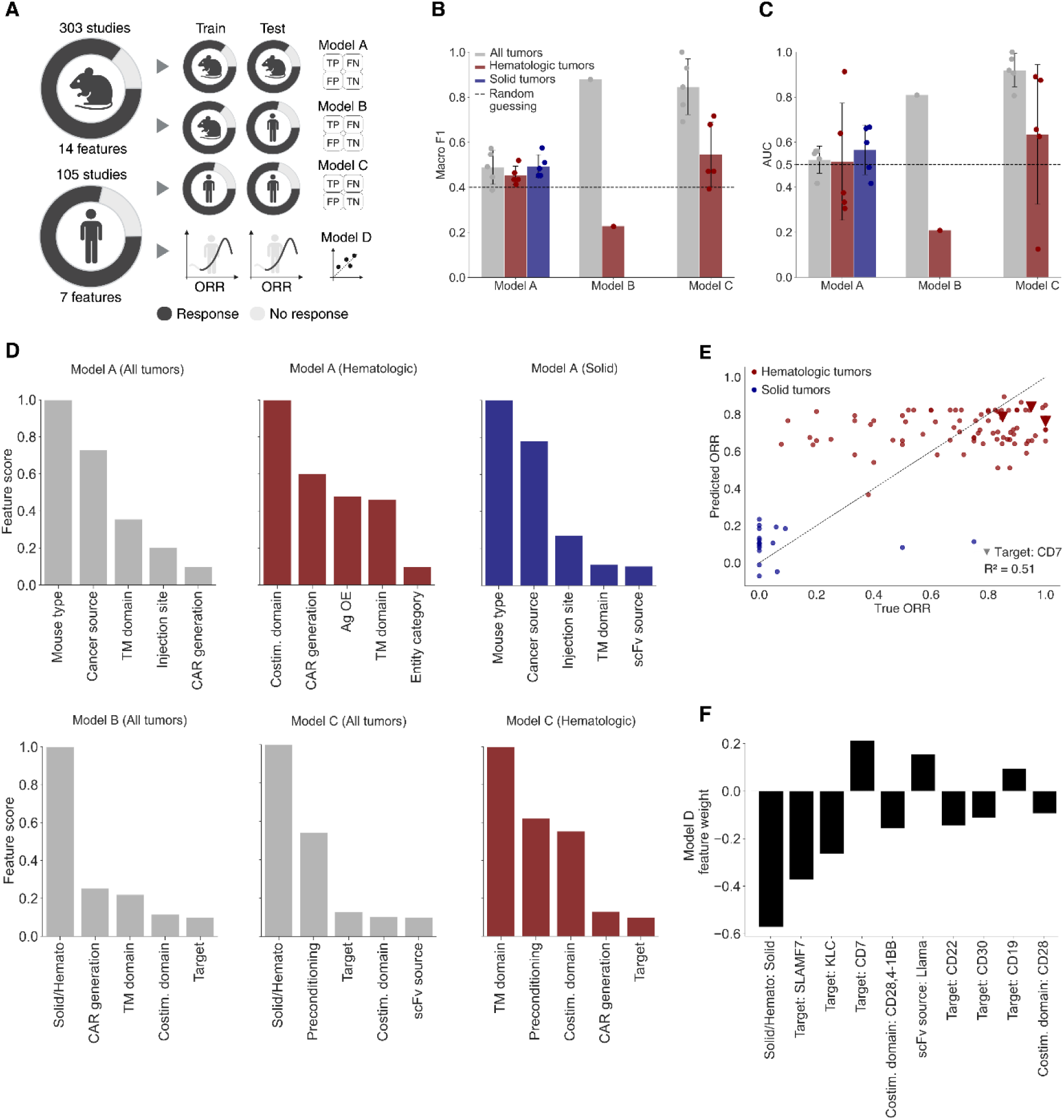
Machine learning analysis of preclinical and clinical datasets identifies tumor type as the most predictive feature across both classification and regression tasks. (A) Schematic outline of the model training and testing strategies: Models A, B, and C are classification models, whereas Model D is a regression model. Models A, C, and D were trained and validated using 5-fold cross-validation; specifically, Model A utilized preclinical data, while Models C and D used clinical data. Model B was trained and validated on preclinical data and subsequently tested on clinical data. (B-C) Performance metrics, including macro F1 score (B) and AUC (C) are reported for Models A, B, and C across the entire dataset (“All tumors”), hematological and solid tumor subsets. Results for the solid tumor subset are only presented for Model A due to the limited size and label imbalance of the clinical solid tumor subset (Supplementary methods). Horizontal, dashed lines indicate the performance of a model using random guessing as a baseline (Supplementary methods). Error bars represent the standard deviation of the scores across the 5 validation folds (shown as individual points) for models A and C. (D) Feature importance scores from Model A (top) and Models B and C ("All tumors") and Model C Hematologic (bottom). Tumor type (solid vs. hematologic) consistently emerges as the most predictive feature for both Models B and C across the entire dataset (‘All tumors’). In Model B, this is followed by ‘CAR generation’, whereas in Model C, ‘Preconditioning’ ranks second. When Model C is trained and validated specifically on the hematologic tumor subset, ‘TM Domain’ becomes the most predictive feature, with ‘Preconditioning’ and ‘Costimulatory domain’ ranking next. All feature importance scores are normalized and expressed on a relative scale. (E) Scatter plots show true versus predicted Overall Response Rates (ORR) across all cross-validation folds for Model D, with a moderate correlation (mean R² = 0.51). (F) Feature weights indicate that lower predicted ORRs are mainly linked to solid tumors, which have the most negative weight.

## Discussion

The translational value of animal models remains a longstanding concern, encompassing more areas than just the CAR-T cell field^46-49^. Preclinical models often fail to accurately replicate human malignancy and the complexity of the immune system, yielding artificial and unreliable results when translated into the clinic^50,51^. The question of how accurately animal models can reflect and predict clinical CAR-T cell outcomes remains highly relevant.

To address the question, a comprehensive database of publicly available CAR-T cell data was compiled, including 105 clinical trials with 3,496 patients. As expected, hematological tumors showed the highest clinical responses, as well as higher rates of CAR-associated AE and hematotoxicity compared to solid malignancies, although severe AE have been observed in CAR-treated patients for both tumor types^52,53^. The majority of the analyzed clinical trials employed anti-CD19 and anti-BCMA CAR-T cells. Besides CD19, BCMA is the only other CAR-T cell target specifically approved by the FDA for the treatment of multiple myeloma^54,55^. Solid tumors accounted for a broader range of targets, accounting for the anticipated heterogeneity of their surface antigen^56-58^. HER2, GD2 and mesothelin appeared as the most frequently investigated solid clinical targets. The prevalence of CD19 and BCMA, coupled with the significantly lower numbers of studies for solid tumors, inevitably skewed the overall trend for efficacy and safety. As more trials with novel solid tumor targets are published, a clearer efficacy assessment across different antigens will emerge.

Our preclinical analysis included 303 experiments, with a total of 2432 mice. CD19, mesothelin, and HER2 were the predominant CAR antigens but CD19 targeting in a B-ALL tumor model represented again a strong bias in the whole analysis. Similarly to the clinical setting, preclinical studies accounted for a small number of solid tumor models with a broad range of different targets. Regardless of the nature of the tumor investigated, the majority of CAR-T cell therapies for blood-borne and solid malignancies were composed of murine scFv domains and either 4-1BB or CD28 costimulatory domains, in line with the first FDA-approved CAR therapies^59^. These figures are not truly representative of the current pipeline for CAR-T cell therapy development, which reports a growing body of trials (27 %) focusing on solid tumors of different origins^60^. Irrespective of the CAR molecule employed, both tumor types reported partial responses in mice, with no advantage for blood-borne malignancies.

Three logistic regression models were trained to predict clinical response from preclinical (B) or clinical (C) data, and preclinical response from preclinical data (A). Given the inherent differences between the efficacy readout of clinical and preclinical studies, we binarized the outcomes into responders and non-responders. This underscored a critical distinction: while the reporting of clinical results is highly standardized, preclinical models rely on surrogate endpoints which may have diverse biological implications. Although necessary for our model, this approach could reduce the granularity of our dataset and overlook subtle patterns in the responses. When predicting clinical response using both clinical and preclinical data, our logistic regression models exhibited superior performance compared to a random classifier. Notably, the predictive strength of these models is predominantly derived from tumor type information. Despite this, the results indicate a degree of concordance between preclinical and clinical studies, even in light of the well-established limitations of animal models in accurately forecasting clinical outcomes in oncology^61-63^. The model predicting preclinical outcomes from preclinical data including solid and hematological tumors (Model A - “All tumors”) showed an overall poor performance (Macro F1 = 0.49 ± 0.07, AUC = 0.52 ± 0.06), indicating that the available data does not capture the complexity of treatment effects in preclinical experimentation. The most discriminating factor for both models predicting clinical responses (B and C) was the cancer’s solid or hematological origin, in line with the reported greater efficacy of CAR-T cells in hematological malignancies compared to solid tumors^64,65^. After subsetting the datasets into solid and hematological tumors, performance of Model B and C dropped considerably, from 0.86 to 0.22 and from 0.85 to 0.55 of Macro F1, respectively. This reduction in performance may be due to the absence of the main discriminating feature, the smaller dataset size, and the increased severity of label imbalance. Consistent with the feature rankings from the classification model, the regression model D assigned the highest negative weight to the "Solid" tumor type. Given a categorical Lasso regression, where a negative weight indicates a decrease in the target variable and its magnitude reflects the strength of such decrease, our findings confirm that solid tumors are linked to overall poorer response rates.

A number of confounding factors should be taken into consideration for this study. Firstly, preclinical research is biased towards positive data, as negative results are rarely published in prestigious journals, unlike in clinical research^66^. Researchers may also employ mouse models to reinforce their hypotheses, by employing control groups in a way that would negatively bias the experimental setup^67^. It was therefore highly challenging to obtain an accurate and comprehensive understanding of preclinical testing from the currently available data.

Secondly, preclinical evaluation of CAR-T cells relies predominantly on immunodeficient xenograft models^25,38^. However, tumor xenografts fail to investigate the impact of endogenous immunity on tumor control, reduce tumor heterogeneity, and belittle toxicity against antigen-expressing healthy tissues^35,41^. As a result, the overall rate of toxicity reporting was little to none for both hematological (6.8 %) and solid tumors (8.2 %), suggesting it to be underreported or not conducted at all. The higher efficacy and lower safety concerns of immunodeficient mouse models make them the most commonly used in vivo models in the field, despite their high maintenance costs^35,50^. This is clearly reflected in our data, whereby immunodeficient and immunocompromised models displayed partial responses on average, in stark contrast to immunocompetent models. The limited information on toxicities prevented any definitive conclusion regarding toxicity profiles in hematological versus solid tumors, highlighting the limitations of preclinical mouse models in accurately predicting clinical toxicity.

Researchers possess a number of potentially more suitable animal models for in vivo preclinical evaluation of immunotherapies, namely syngeneic and humanized models^68^. Unfortunately, fully syngeneic mouse models constituted a very small proportion (4 %) of overall preclinical records. Such scarcity, combined with insufficient toxicity reporting in preclinical publications, highlighted the strong bias towards immunodeficient models^38^. Considering these low numbers and the phenomenon of positive publication bias, it is challenging to assess whether immunocompetent mice would be better predictors of clinical response. Bearing in mind the key importance of immunity in immunotherapies, it is tempting to speculate that fully immunocompetent models may better discriminate towards clinical outcome. This hypothesis will require adequate investigation and demonstration.

Thirdly, this work did not account for a number of factors like the CAR transduction vector, the in vitro handling and expansion of cancer and CAR-T cells prior to animal treatment, the differences in dosing regimens, and the timing of in vivo treatment. Furthermore, the strict exclusion criteria here applied led to the removal of CAR therapies involving immune cells other than T lymphocytes and the combination of CAR-T cells with other strategies like antibodies, cytokines, or costimulatory receptors. The use of combination therapies involving CAR-T cells is an ever-growing field of research^69^. In this analysis, we observed a strong interest in exploiting the heterogenous range of identified solid tumor antigens and can thus anticipate a broader exploration of more sophisticated CAR therapy designs, multi-antigen targeting strategies and combinatorial approaches to overcome the challenges posed by solid malignancies^70,21,71,72^.

In conclusion, while our machine learning models indicated weak predictive power of clinical responses from preclinical data, the findings underscored the need for more diverse and comprehensive preclinical studies. The integration of more immunocompetent and humanized mouse models, with more standardized testing and reporting guidelines, will allow a more robust analysis of the weight and predictive power of each feature. Such an approach will hopefully lead to more effective and tailored CAR-T cell therapies for both hematological and solid tumors.

## Supplementary Figures

**Supplementary Figure 1:**
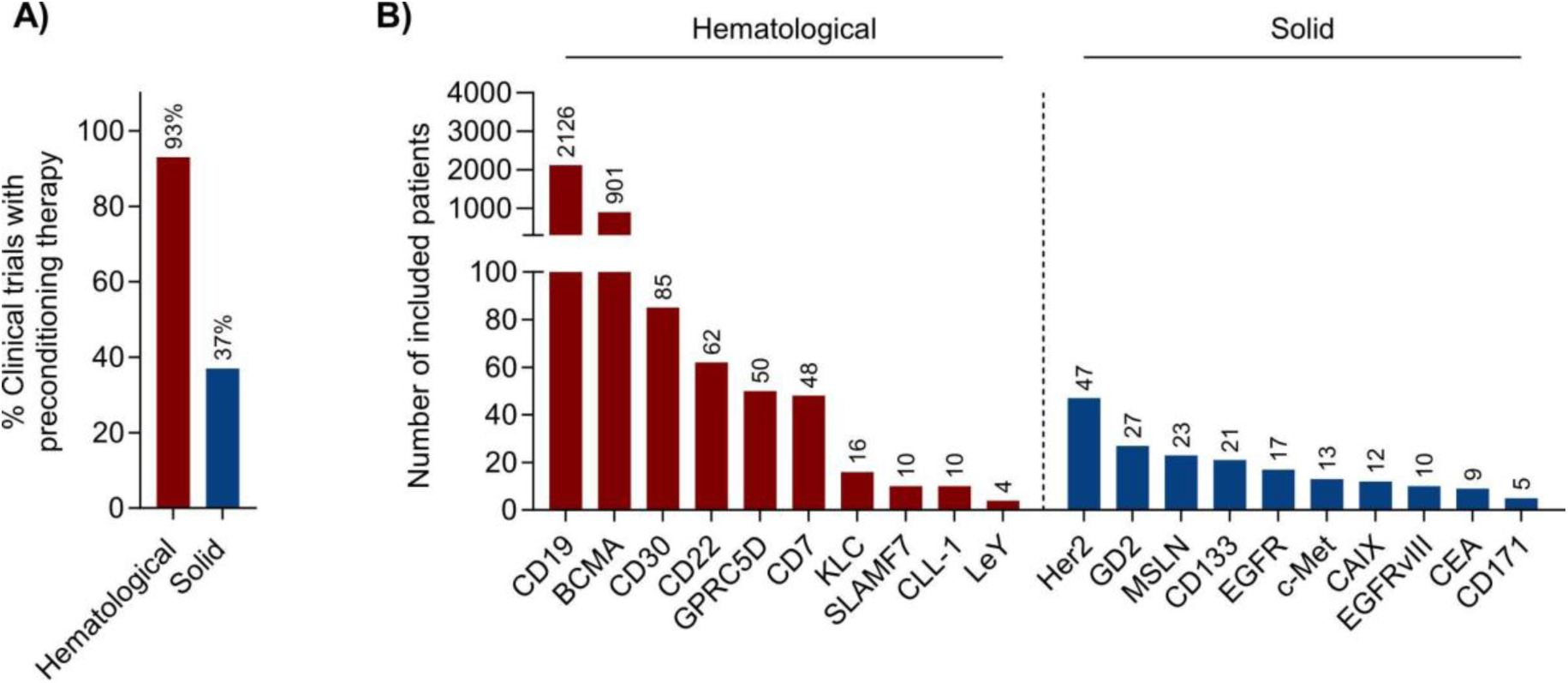
Safety data and treatment responses for hematological and solid clinical trials of CAR-T cells. (A) Implementation of preconditioning therapy, in terms of chemo- or radiation therapy. (B) Distribution of studied patients by target antigen.

**Supplementary Figure 2:**
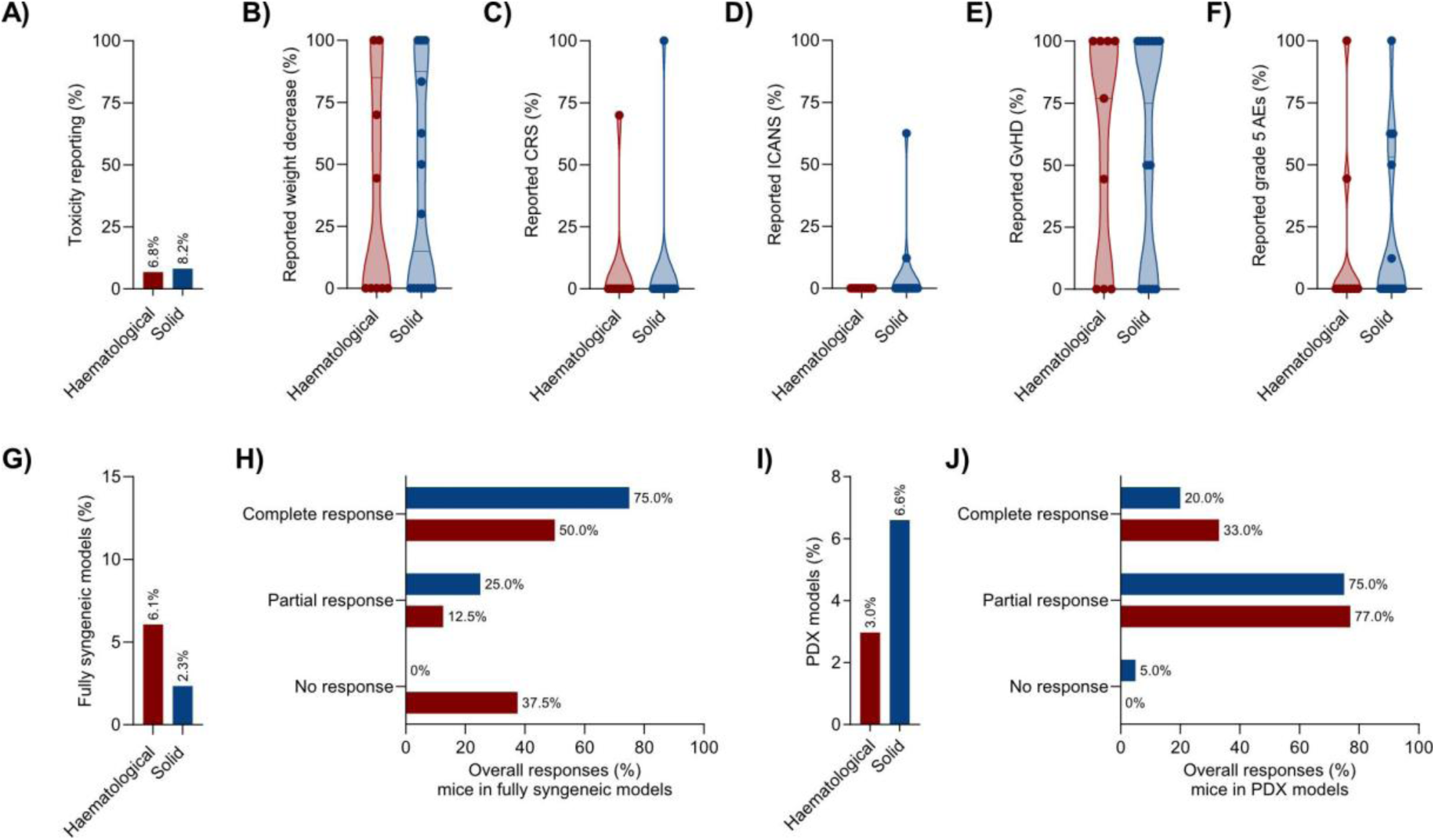
Safety evaluation and treatment responses for hematological and solid preclinical studies of CAR-T cells. (A) Rates of toxicity reporting for CAR-T cell therapy in hematological and solid tumor models. (B) Proportions of toxic events in experimental animals such as weight loss (B), CRS (C), ICANS (D), GvHD (E) and lethal AE (F) for either hematological or solid tumors. (G) Sum of all immunocompetent mice employed in fully syngeneic tumor models belonging to CAR treatment groups for either category of tumors. (H) Overall responses as for tumor clearance, decrease, slower growth or progression for fully syngeneic mouse models. (I) Sum of all PDX models implemented in preclinical studies. (J) Response distribution across all PDX models used.

## Data Sharing Statement

The raw data presented in this study may be found in a data supplement (extended data file 1) available with the online version of this article. For original data or further inquiries, please contact

## Acknowledgements

This study was supported by the Bavarian Cancer Research Center (BZKF) (TANGO to S.K.), the Deutsche Forschungsgemeinschaft (DFG, KO5055-2-1 and KO5055/3-1 to S.K.), the international doctoral program ‘i-Target: immunotargeting of cancer’ (funded by the Elite Network of Bavaria; to S. K.), the Melanoma Research Alliance (grant number 409510 to S.K.), Marie Sklodowska-Curie Training Network for Optimizing Adoptive T Cell Therapy of Cancer (funded by the Horizon 2020 programme of the European Union; grant 955575 to S.K.), Else Kröner-Fresenius-Stiftung (IOLIN to S.K.), German Cancer Aid (AvantCAR.de to S.K.), the Wilhelm-Sander-Stiftung (to S. K.), Ernst Jung Stiftung (to S.K.), Institutional Strategy LMUexcellent of LMU Munich (within the framework of the German Excellence Initiative; to S.K.), the Go-Bio-Initiative (to S.K.), the m4-Award of the Bavarian Ministry for Economical Affairs (to S.K.), Bundesministerium für Bildung und Forschung (to S.K.), European Research Council (Starting Grant 756017, PoC Grant 101100460 and CoG 101124203 to S. K.), by the SFB-TRR 338/1 2021–452881907 (to S.K.), Fritz-Bender Foundation (to S.K.), Deutsche José Carreras Leukämie Stiftung (to S.K.), Hector Foundation (to S.K.), Bavarian Research Foundation (BAYCELLATOR to S.K.), the Bruno and Helene Jöster Foundation (360° CAR to S.K) and the Monika-Kutzner Foundation (to S.K.). L.G. received funding from the Studienstiftung des Deutschen Volkes. Figures were created with BioRender.com.

## Authorship contributions

Conceptualization: D.A.S., L.G., E.C., C.M., S.K. Investigation: D.A.S, L.G., E.C., D.S. Formal analysis: D.A.S., L.G., E.C., D.S, C.M., S.K. Supervision: C.M., S.K. Writing: D.A.S., L.G., E.C., D.S., S.K. All authors contributed to feedback and proofreading.

## Data Availability

The main data supporting the results in this study are available within the paper and its Supplementary Information. The raw and analyzed datasets generated during the study are available for research purposes from the corresponding authors on reasonable request.

## Informed consent

All authors approve the manuscript for publication.

## Supplementary Material

Refer to the web version for supplementary material.

## Supplementary Methods

### Information sources, search strategy and data collection process

The clinical trial records were sourced from PubMed and ClinicalTrials.gov until December 1st, 2023, employing specific search criteria. The results from PubMed were retrieved using the query "((CAR-T) OR (Chimeric antigen receptor)) AND (Clinical Trial[Publication Type]) AND (English[Language])". Similarly, in ClinicalTrials.gov, the search query “(CAR-T) OR (Chimeric antigen receptor)” was used in the “other terms” search field, with an additional filter for completed and terminated trials to allow for the selection of studies with reported results. To streamline the dataset, papers obtained from PubMed underwent manual screening to extract the clinical trial identifiers. For the clinical trial identifiers with NCT format, the records were retrieved from ClinicalTrials.gov and duplicates were removed. For the clinical trials with other formats, the information was manually retrieved from the relevant sources, if available.

The preclinical records were obtained from PubMed until December 1st, 2023, using the following search: “((CAR-T) OR (chimeric antigen receptor)) AND ((CD19) OR (TNFRSF17) OR (CD269) OR (BCMA) OR (Siglec-2) OR (CD22) OR (Mesothelin) OR (Her2) OR (ERBB2) OR (CD30) OR (TNFRSF8) OR (MSLN) OR (GD2) OR (CD7) OR (ERBB1) OR (CEA) OR (CEACAM) OR (SLAMF7) OR (CD319) OR (CS1) OR (LeY) OR (KLC) OR (EGFRvIII) OR (EGFR) OR (CLL-1) OR (CE7) OR (L1CAM) OR (CD171) OR (AC133) OR (CD133) OR (CA9) OR (CAIX) OR (HGFR) OR (c-Met)) NOT (Review[Publication Type]) NOT (Clinical trial[Publication Type]) NOT (Systematic Review[Publication Type]) NOT (Meta-Analysis[Publication Type]) AND (English[Language])”. These included all publications in English regarding CAR-T cells and the targets from the previously included clinical trials (including their commonly used synonyms), excluding reviews, systematic reviews, meta-analyses and studies of clinical nature.

To prevent biases in the assessment, each entry was evaluated in all its aspects, including inclusion/exclusion criteria and data extraction, by two reviewers independently. Disagreements were resolved by discussion or with the intervention of a third reviewer.

### Further explanation regarding eligibility criteria and selection process

An additional clarification of the exclusion criteria considered by the investigators as the most interpretative is provided: the exclusion criterion ‘Combination therapy’ refers to studies in which the CAR-T cell therapeutic was not the sole object of investigation for efficacy in the clinical and preclinical settings, apart from chemo- and radiotherapy. This included, among others, knock-out and/or knock-in mutations, monoclonal antibodies, bispecific CAR molecules and chimeric switch receptors.

Studies were classified as ‘Not about CAR efficacy’ in the event of studies investigating, for example, the safety profile of certain suicide switches, novel methods of CAR generation and ex vivo expansion.

Studies were classified as ‘Non-classical CAR setting’ where the main study subject was a CAR molecule modified in its classical structure and mode of function. Examples include, but are not limited to, VHH-, nanobody-, or NKG2D-based CAR platforms, SynNotch and logic-gated systems.

Studies were excluded for ‘Incomplete data’ if they did not provide relevant information about the full CAR structure and/or the efficacy of the treatment. Preclinical work deemed to be ‘Studying irrelevant variables in the experimental set-up’ was characterized by an effort to elucidate other aspects of the adoptive use of CAR-T cells other than efficacy in animal models, examples being naïve vs effector phenotypes, higher affinity binding moieties, mapping of intracellular phosphorylation patterns or the composition of the TME.

### Machine learning-guided analysis of our database

#### Data collection and preprocessing

The input data for the Machine Learning analysis comprised both preclinical and clinical CAR-T cell studies. The preclinical dataset consisted of 303 data points, each representing a distinct study with features such as ‘Target’, ‘Solid or Hematologic tumors’, ‘Entity category’, ‘Cancer source’, ‘Antigen origin’, ‘Ag OE’, ‘Injection site’, ‘Mouse type’, ‘CAR generation’, ‘scFv source’, ‘TM domain’, ‘Costimulatory domain’, ‘Preconditioning’, and ‘Number of Mice’ (Extended data file 1). The response variable for the preclinical dataset was initially categorized into three classes (’No response’, ‘Partial response’, ‘Complete response’) and subsequently binarized into ‘No response’ (negative class) and a combined positive class comprising the other two categories to train the logistic regression models.

The clinical dataset consisted of 105 data points with features including ‘Target’, ‘Solid or Hematologic tumors’, ‘Entity category’, ‘CAR generation’, ‘scFv source’, ‘TM domain’, ‘Costimulatory domain’, ‘Preconditioning’, and ‘Number of Participants’ (Table 2 and 3).

Out of the initial set of features, ‘Entity category’ was removed, since several clinical trials include multiple entities in the same study. ‘Number of Mice’ and ‘Number of Participants’ were also neglected, as they are not expected to have an effect on the outcome of the trial or preclinical study.

The response variable ORR, originally a continuous variable ranging from 0 to 1, was binarized using a 0.25 cutoff. The ‘Preconditioning’ variable was also binarized to indicate the presence or absence of preconditioning.

#### Model training and hyperparameter optimization

##### Classification Models

1. Model A: Preclinical data training and validation

● Data Preparation: The preclinical dataset was used to train the model. Features were one-hot encoded and used to form the final training set.
● Model Definition: A logistic regression model with objective function:

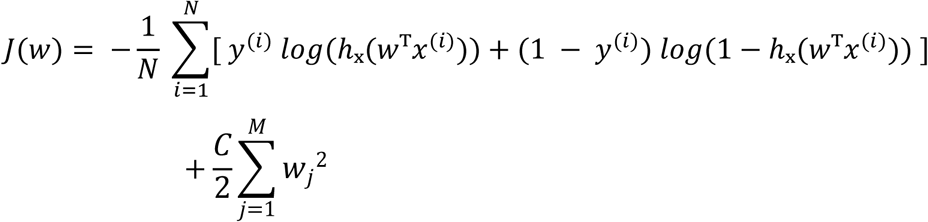

Where:

-*N* is the number of training examples.
-*M* is the number of training parameters.
-*y*^(*i*)^ is the true label (0 or 1) for the *i*-th training example.
-*x*^(*i*)^ is the feature vector for the *i*-th training example.
-*w* is the weight vector (parameters).
-ℎₓ(*w^T^x*^(*i*)^) = 1 / (1 + *exp*(*w^T^x*^(*i*)^)) is the logistic (sigmoid) function.
-*C* is the regularization strength, which controls the trade-off between fitting the data and penalizing large model parameters to prevent overfitting. A higher value of *C* increases the penalty on large weights, making the model simpler and less prone to overfitting. The following were defined as hyperparameters to tune:

○ *C*: Regularization strength, values ranged from 1e-4 to 1e4
○ *penalty*: ‘l2’
○ *solver*: [’lbfgs’, ‘newton-cg’, ‘newton-cholesky’] Class weights were balanced to account for label imbalance.
● Hyperparameter Optimization: A grid search over the defined parameter space with 5-fold cross-validation was performed. Multiple scoring metrics were used, including Area Under the Curve (AUC), Macro F1, accuracy, and average precision. The final model was selected based on the best average precision score.
● Model Training and Validation: The model was trained and validated using a 5-fold cross-validation strategy on the preclinical dataset. This approach ensured that the model’s performance was assessed on multiple subsets of the data, providing a more robust evaluation of its generalizability.
2. Model B: Preclinical data training and clinical data testing

● Data Preparation: The model was trained on the preclinical dataset and tested on the clinical dataset. Only features common to both datasets (’’Solid or Hematologic tumors’’, ‘scFv source’, ‘Target’, ‘CAR generation’, ‘TM domain’, ‘Preconditioning’, ‘Costimulatory domain’) were used to ensure compatibility. The same preprocessing steps as in Model A were applied.
● Model Definition and Hyperparameter Optimization: The logistic regression model was defined, and hyperparameters optimized as described for Model A. The optimal hyperparameters identified were used to train the model on the preclinical data.
● Model Testing: The trained model was then tested on the clinical dataset to evaluate its predictive performance on external data. Performance metrics such as the Area Under the Receiver Operating Characteristic Curve (ROC AUC), Macro F1, accuracy, and average precision were calculated.
3. Model C: Clinical data training and validation

● Data Preparation: The clinical dataset was used for both training and validation. The same preprocessing steps as in Model A were applied to this dataset.
● Model Definition and Hyperparameter Optimization: The logistic regression model was defined, and hyperparameters optimized as described fo*r Model A. The* optimal hyperpa*ramet*e*rs* identified were used to train the model on the clinical data.
● Model Training and Validation: Similar to Model A, a 5-fold cross-validation strategy was employed to train and validate the model on the clinical dataset.
4. Random model

● A random baseline model, assigning response labels (“Non-responders” and “Responders”) with a 50% probability for each, was implemented for each data subset. These subsets included preclinical data categories: “All tumors,” “Hematologic,” and “Solid”; and clinical data categories: “All tumors” and “Hematologic.” This model served as a benchmark for evaluating classification metrics.

For each model, performance metrics AUC, macro F1 score, sensitivity, and recall were calculated. The AUC reflects the model’s ability to distinguish between positive (“responders”) and negative (“non-responders”) classes, with higher values indicating better performance. The Macro F1 score, a class-agnostic metric, calculates the harmonic mean of precision and recall, making it particularly useful for imbalanced class distributions. Higher macro F1 values indicate superior performance, demonstrating the models’ effectiveness in handling class imbalances in treatment responses. Sensitivity measures the proportion of actual responders correctly identified by the model, and specificity measures the proportion of true non-responders accurately classified.

Feature importance was determined by extracting coefficients from a trained logistic regression model. The coefficients for each feature were computed across all response classes, and the absolute values of these coefficients were aggregated by the original feature names. This process allowed for the identification of the most influential features contributing to the model’s predictions.

#### Subset Analysis

Both preclinical and clinical datasets were divided into hematological and solid tumors:

● Preclinical: 171 solid tumor data points, 132 hematological tumor data points
● Clinical: 86 hematological tumor data points, 19 solid tumor data points For each subset, models A, B, and C were trained, and the best hyperparameters were identified. The performance metrics and feature importance rankings were computed and reported for the best-performing models.

#### Regression Model

Model D, a L1-regularized linear regression model (Lasso regression) was trained on the clinical dataset to predict the continuous ORR. The training pipeline consisted of the following steps:

● Preprocessing: Categorical variables were encoded using one-hot encoding.
● Model Definition: A linear regression model with objective function:

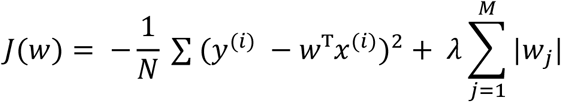

Where:

-*J*(*w*) is the objective function.
-*N* is the number of training examples.
-*M* is the number of training parameters.
-*y*^(*i*)^ is the true value for the *i*-th training example.
-*x*^(*i*)^ is the feature vector for the *i*-th training example.
-*w* is the weight vector.
-λ is the regularization parameter that controls the penalty’s strength. The following were defined as hyperparameters to tune:

○ λ: Regularization strength, values ranged from 1e-4 to 1e4
● Model Training: A Lasso regression model was trained, with hyperparameter optimization performed using a grid search to identify the optimal alpha parameter. The best model was selected based on cross-validation performance, with significant features identified by their non-zero coefficients.
● Evaluation: The model’s performance was assessed using the coefficient of determination (R²), calculated from the relationship between the true and predicted ORR values for each cross-validation fold.
● Feature Analysis: Non-zero feature weights from the Lasso regression were extracted and ranked by their absolute magnitude to determine their relative importance in predicting the ORR.

#### Environment

The analysis was conducted using a conda environment with Python 3.10.13, scikit-learn 1.3.2, and seaborn 0.13.0 for plotting.

The code used for the machine learning analysis is available at https://github.com/DanScarc/CAR-T-Meta-Analysis.git.

